# Effect modification of *FADS2* polymorphisms on the association between breastfeeding and intelligence: results from a collaborative meta-analysis

**DOI:** 10.1101/184234

**Authors:** Fernando Pires Hartwig, Neil Martin Davies, Bernardo Lessa Horta, Tarunveer S. Ahluwalia, Hans Bisgaard, Klaus Bønnelykke, Avshalom Caspi, Terrie E. Moffitt, Richie Poulton, Ayesha Sajjad, Henning W Tiemeier, Albert Dalmau Bueno, Mònica Guxens, Mariona Bustamante Pineda, Loreto Santa-Marina, Nadine Parker, Tomáš Paus, Zdenka Pausova, Lotte Lauritzen, Theresia M. Schnurr, Kim F. Michaelsen, Torben Hansen, Wendy Oddy, Craig E. Pennell, Nicole M. Warrington, George Davey Smith, Cesar Gomes Victora

**Affiliations:** Postgraduate Program in Epidemiology, Federal University of Pelotas, Pelotas, Brazil; Medical Research Council Integrative Epidemiology Unit, University of Bristol, Bristol, United Kingdom; School of Social and Community Medicine, University of Bristol, Bristol, United Kingdom; COPSAC, Copenhagen Prospective Studies on Asthma in Childhood, Herlev and Gentofte Hospital, Faculty of Health Sciences, University of Copenhagen, Copenhagen, Denmark; Duke University, Durham, USA; Institute of Psychiatry, Psychology, and Neuroscience, King’s College London, London, United Kingdom; Department of Psychology, University of Otago, Dunedin, New Zealand; Department of Epidemiology, Erasmus University Medical Centre, Rotterdam, The Netherlands; Department of Child and Adolescent Psychiatry/Psychology, Erasmus University Medical Centre, Rotterdam, The Netherlands; ISGlobal, Centre for Research in Environmental Epidemiology (CREAL), Barcelona, Spain; Universitat Pompeu Fabra (UPF), Barcelona, Spain; CIBER Epidemiología y Salud Pública (CIBERESP), Madrid, Spain; Centre for Genomic Regulation (CRG), The Barcelona Institute of Science and Technology, Barcelona, Spain; BIODONOSTIA Health Research Institute, San Sebastian, Spain; Public Health Division of Gipuzkoa, San Sebastian, Spain; Rotman Research Institute, Baycrest Centre for Geriatric Care, Toronto, Canada; Institute of Medical Science, University of Toronto, Toronto, Canada; Departments of Psychiatry and Psychology, University of Toronto, Toronto, Canada; Hospital for Sick Children Research Institute, Peter Gilgan Centre for Research and Learning, Toronto, Canada; Department of Nutritional Sciences, University of Toronto, Toronto, Canada; Department of Physiology, University of Toronto, Toronto, Canada; Department of Nutrition, Exercise and Sports, Faculty of Science, University of Copenhagen, Copenhagen, Denmark; Novo Nordisk Foundation Centre for Basic Metabolic Research, Section of Metabolic Genetics, Faculty of Health and Medical Sciences, University of Copenhagen, Copenhagen, Denmark; Menzies Institute for Medical Research, University of Tasmania, Hobart, Australia; School of Women’s and Infants’ Health, The University of Western Australia, Perth, Australia; The University of Queensland Diamantina Institute, The University of Queensland, Translational Research Institute, Brisbane, Australia

**Keywords:** Breastfeeding, Intelligence, *FADS2*, Fatty acids, Effect modification, Meta-analysis.

## Abstract

**Background:** Accumulating evidence suggests that breastfeeding benefits the children’s intelligence. Long-chain polyunsaturated fatty acids (LC-PUFAs) present in breast milk may explain part of this association. Under a nutritional adequacy hypothesis, an interaction between breastfeeding and genetic variants associated with endogenous LC-PUFAs synthesis might be expected. However, the literature on this topic is controversial.

**Methods and Findings:** We investigated this Gene×Environment interaction in a *de novo* meta-analysis involving >12,000 individuals in the primary analysis, and >45,000 individuals in a secondary analysis using relaxed inclusion criteria. Our primary analysis used ever breastfeeding, *FADS2* polymorphisms rs174575 and rs1535 coded assuming a recessive effect of the G allele, and intelligence quotient (IQ) in Z scores. Using random effects meta-analysis, ever breastfeeding was associated with 0.17 (95% CI: 0.03; 0.32) higher Z scores in IQ, or about 2.1 points. There was no strong evidence of interaction, with pooled covariate-adjusted interaction coefficients (i.e., difference between genetic groups of the difference in IQZ scores comparing ever with never breastfed individuals) of 0.12 (95% CI: −0.19; 0.43) and 0.06 (95% CI: −0.16; 0.27) for the rs174575 and rs1535 variants, respectively. Secondary analyses corroborated these results. In studies with ≥5.85 and <5.85 months of breastfeeding duration, pooled estimates for the rs174575 variant were 0.50 (95% CI: −0.06; 1.06) and 0.14 (95% CI: −0.10; 0.38), respectively, and 0.27 (95% CI: −0.28; 0.82) and −0.01 (95% CI: −0.19; 0.16) for the rs1535 variant. However, between-group comparisons were underpowered.

**Conclusions:** Our findings do not support an interaction between ever breastfeeding and *FADS2* polymorphisms. However, our subgroup analysis raises the possibility that breastfeeding supplies LC-PUFAs requirements for cognitive development (if such threshold exists) if it lasts for some (currently unknown) time. Future studies in large individual-level datasets would allow properly powered subgroup analyses and would improve our understanding on the role of breastfeeding duration in the breastfeeding×*FADS2* interaction.

## Introduction

Breastfeeding has well-established short term benefits on children’s health. There is also accumulating evidence that breastfeeding may also benefit cognitive development^1^. A recent meta-analysis of observational studies reported that breastfed subjects scored higher on intelligence quotient (IQ) tests [mean difference 3.4 (95% CI: 2.3; 4.6)] than non-breastfed subjects^2^. Although issues such as residual confounding^3^ and publication bias^4^ may have affected this estimate, randomised controlled trials of breastfeeding promotion reported benefits in motor development in the first year of life^5^ and in IQ at 6.5 years of age^6^. Additional studies corroborate the notion that breastfeeding has a causal effect on IQ. These include comparisons between cohorts with different confounding structures^7^, and between mothers who tried, but could not breastfeed their child, and mothers who had formula feeding as their first choice^8^.

One of the possible biological mechanisms underlying the effect of breastfeeding on IQ is through long-chain polyunsaturated fatty acids (LC-PUFAs), such as docosahexaenoic acid (DHA). Meta-analyses of randomised controlled trials of supplementation of DHA and other LC-PUFAs in infants reported improved cognitive development^9^ and visual acuity^10^. Indeed, DHA is an important component of the membrane of brain cells and retina cells^11,12^. Studies in animal models and humans suggest that adequate levels of DHA are important for cognitive development through influencing several processes, such as biogenesis and fluidity of cellular membranes, neurogenesis, neurotransmission and protection against oxidative stress^12,13^.

The role of LC-PUFAs in the association between breastfeeding and IQ can be investigated through a Gene×Environment (G×E) interaction analysis. For example, it is possible that there is an upper limit for the benefits of increasing DHA levels and such requirements are met by pre-formed DHA available in breast milk. In this case, inter-individual variation in IQ due to genetically determined differences in DHA endogenous synthesis from metabolic precursors would only be observable in individuals who were not breastfed^14^. This G×E interaction has been investigated using single nucleotide polymorphisms (SNPs) in the *FADS2* gene^14–18^. This gene encodes a desaturase enzyme that catalyses a rate-limiting reaction in the LC-PUFAs pathway^19,20^. Candidate gene and genome-wide approaches reported that minor alleles of SNPs in the *FADS2* gene were associated with lower levels of PUFAs in plasma and erythrocyte phospholipids^21-24^.

Caspi et al. were the first to evaluate the interaction between genetic variation in *FADS2* and breastfeeding, with IQ in children as the outcome. Two SNPs were evaluated: rs1535 (major/minor alleles: A/G) and rs174575 (major/minor alleles: C/G). For both SNPs, having ever being breastfed was positively associated with IQ in all genetic groups, except in G-allele homozygotes, where there was no association^15^. Although there was evidence for a GxE interaction, it was not consistent with the nutritional adequacy hypothesis outlined above. However, in a replication study, Steer et al. results were consistent with the nutritional adequacy hypothesis (and therefore inconsistent with Caspi et al.’s findings), with breastfed individuals presenting similar mean values of IQ across *FADS2* genotypes. Such values were higher than those observed in never breastfed individuals, with the lowest value (and thus the greatest effect of breastfeeding) being in GG individuals^14^. Morales et al. reported that a negative association between genotypes in other genetic variants related to lower activity of enzymes involved in elongation and desaturation processes and cognition was only evident in non-breastfed individuals^25^. However, three studies in twins (but not twin studies, in the sense that they did not aim at estimating heritability) did not detect strong evidence supporting this G×E interaction^16-18^.

In this study, we aimed at improving the current understanding on this G×E interaction and gaining insights into the sources of heterogeneity between studies through a consortium-based initiative^26^.

## Methods

### Overview of the study protocol

The protocol of this study has been published elsewhere^26^. Briefly, studies that were known by the coordinating team to have at least some of the data required available, as well as other studies suggested by collaborators, were invited to participate. All studies that were contacted (and were eligible) accepted to participate.

All of the following criteria were required for eligibility: i) availability of at least a binary breastfeeding variable (i.e., whether or not the study individuals where ever breastfed), intelligence measured using standard tests, and at least rs174575 or rs1535 SNPs (either genotyped or imputed); and ii) European-ancestry studies, or multi-ethnic studies if possible to define a subsample of European ancestry individuals. Exclusion criteria were: i) only poorly imputed genetic data were available (metrics of imputation such as *r*^2^ or INFO quality below 0.3); ii) twin studies; iii) lack of appropriate ethical approval.

Data analysis was performed locally by data analysts of the collaborating studies. Standardised analysis scripts written in R (http://www.r-project.org/) were prepared centrally and distributed to the analysts, along with a detailed analysis plan and instructions to format the data. The scripts automatically generated files containing summary descriptive and association statistics, which were centrally meta-analysed.

As the analyses progressed, some modifications in the original protocol were required. These are described in the Supplementary Methods.

### Participating studies

A total of 10 eligible studies were identified, all of which were included in the meta-analysis: the 1982 Pelotas Birth Cohort Study^27,28^, Dunedin Multidisciplinary Health and Development Study^15^, Avon Longitudinal Study of Parents and Children (ALSPAC)^29^, Copenhagen Prospective Study on Asthma in Childhood (COPSAC) 2010^30,31^, Generation R Study^32-34^, *INfancia y Medio Ambiente* (INMA) Project^35^, Western Australian Pregnancy Cohort (Raine) Study^36-38^, *Småbørn Kost Og Trivsel*-1 (SKOT-I)^39,40^, SKOT-II^41,42^ and Saguenay Youth Study (SYS)^43,44^.

In addition, a subsample of 32,842 individuals from the UK Biobank^45^ was included. However, this subsample did not fulfil the pre-established eligibility criteria because IQ was not measured using a standard test. Therefore, these data were used in secondary analyses only.

Information about the participating studies is shown in Supplementary Tables 1-3.

### Statistical analyses

The main outcome variable was IQ. IQ tests varied between studies (Supplementary Table 1), so IQ measures were converted to Z scores (mean=0 and variance=1) within each participating study. The primary analysis involved breastfeeding (coded as never=0 and ever=1), *FADS2* polymorphism assuming a recessive genetic effect of the G allele (i.e., GG individuals=1; heterozygotes and non-G allele homozygotes=0) and an interaction term between them. Different genetic effects, different categorizations of breastfeeding, and exclusive breastfeeding (defined as receiving only breast milk and no other food or drink, including water) were evaluated in pre-planned secondary analyses. Unless explicitly stated, all analyses refer to any quality of breastfeeding (i.e., combining exclusive and non-exclusive).

Three analysis models were performed: (i) unadjusted (i.e., no covariates); (ii) adjusted 1: controlling for sex and age (linear and quadratic terms) when IQ was measured, ancestry-informative principal components (when available) and genotyping centre (for studies involving multiple laboratories); (iii) adjusted 2: same covariates in “adjusted 1” model, as well as maternal education (linear and quadratic terms) and maternal cognition (linear and quadratic terms); if only one of the maternal variables was available, adjusted model 2 controlled only for that variable. Continuous covariates, as well as sex (which was coded as male=0 and female=1), were mean-centred before analysis, and squaring was performed before mean centring. Covariate adjustment was performed by including not only a “main effect” term, but also (*FAD52*×Covariate) and (Breastfeeding×Covariate) interaction terms^46^.

As a sensitivity analysis, the role of gene-environment correlation was evaluated by repeating models i) and ii), but having maternal cognition (in Z scores) or maternal schooling (in years) as outcome variables rather than the participant’s IQ. Maternal cognition or schooling are important predictors of an individual’s IQ, and cannot be consequences of the participant’s genotype. Therefore, any evidence of breastfeeding×*FADS2* interaction in this analysis is indicative that those maternal variables may confound the main breastfeeding×*FADS2* interaction analysis (i.e., having participant’s IQ as the outcome variable).

Analyses were performed using linear regression with heteroskedasticity-robust standard errors. Results from all studies were pooled using fixed and random effects meta-analysis. Random effects meta-regression was used to evaluate the potential moderating role of the following variables: IQ test; adjustment for ancestry-informative principal components; age at IQ measurement; timing of breastfeeding measurement; continental region; mean year of birth; prevalence of having ever being breastfed; mean breastfeeding duration; and sample size.

## Results

### Characteristics of participating studies

As shown in Supplementary Table 1, seven out of the 10 eligible studies were conducted in Europe, four were population-based and two were multi-ethnic. The average year at birth ranged from 1972 to 2011. Three studies measured breastfeeding prospectively, and four measured IQ using the Wechsler Intelligence Scale (two for children and two for adults).

Supplementary Table 2 provides a description of the two *FADS2* SNPs in each study. The SNPs rs174575 and rs1535 were directly genotyped in three and five studies, respectively. The minimum value of imputation quality was 0.984. The frequency of the G allele ranged from 20.5% to 30.8% for the rs174575 variant, and from 28.5% to 39.1% for the rs1535 variant. There was no strong statistical evidence against Hardy-Weinberg Equilibrium, with the smallest P-values being 0.058 (Generation R), 0.074 (SKOTI-II) for rs174575, and 0.085 (1982 Pelotas Birth Cohort), 0.044 (Raine) and 0.089 (SKOTI-II) for rs1535. Although these results may be suggestive of some population substructure (especially in Generation R and in the 1982 Pelotas Birth Cohort, which are multi-ethnic studies) or batch effects (especially in SKOTI-II, which is a combination of two independent studies), it is unlikely that such phenomena substantially influenced the results because ancestry-informative principal components computed using genome-wide genotyping data were available and adjusted for in these four studies.

Additional study characteristics are displayed in Supplementary Table 3. Among eligible studies (i.e., excluding the UK Biobank), the mean age, maternal education, and breastfeeding duration ranged from 2.5 to 30.2 years, 11 to 19 years, and 2.3 to 8.2 months, respectively. All IQ measures produced a variable with mean close to 100 and similar standard deviations (median: 12.2; range: 9.6 to 16.3). The exception was the one used in SKOT-I and SKOT-II (i.e., third edition of the Ages and Stages Questionnaire), which produced a variable with mean close to 50.

### Primary analysis

In analyses without stratification according to genotype, ever breastfeeding was associated with increases of 0.37 (95% CI: 0.32; 0.42) and 0.30 (95% CI: 0.20; 0.40) Z scores in IQ in fixed and random effects meta-analysis, respectively. Assuming that a Z score corresponds to 12.2 points (the median of the standard deviation of IQ measures among participating studies), these coefficients correspond to 4.5 and 3.7 points in IQ. In the fully adjusted model (adjusted 2), the respective coefficients were 0.26 (95% CI: 0.21; 0.32) and 0.17 (95% CI: 0.03; 0.32), or 3.2 and 2.1 points in IQ.

Table 1 and Figure 1 display the results of the primary analysis. There was considerable between-study heterogeneity. Among non-G carries for the rs174575 SNP, pooled random effects estimates of IQ Z scores according to breastfeeding (ever=1; never=0) were 0.29 (95% CI: 0.17; 0.40) and 0.15 (95% CI: 0.00; 0.31) in the unadjusted and fully-adjusted models, respectively. Among GG individuals, the respective estimates were 0.43 (95% CI: 0.16; 0.70) and 0.31 (95% CI: 0.05; 0.58). There was no strong evidence of interaction, with pooled estimates of the breastfeeding×*FADS2* interaction term of 0.18 (95% CI: −0.18; 0.54) and 0.12 (95% CI: −0.19; 0.43), respectively. These coefficients can be interpreted as the difference between genetic groups of the difference in IQ Z scores comparing ever with never breastfed individuals. Similar results were obtained when using fixed effects meta-analysis.

Results for the rs1535 variant were presented a similar trend, but were even less suggestive of interaction. When using random effects meta-analysis, the estimates of the interaction term were −0.04 (95% CI: −0.24; 0.15) and 0.06 (95% CI: −0.16; 0.27) in the unadjusted and fully-adjusted models, respectively. Using fixed effects meta-analysis yielded similar results.

**Table 1.** Meta-analytical linear regression coefficients (β) of cognitive measures (in standard deviation units) according to breastfeeding (never=0; ever=l), within strata of *FADS2* rs174575 or rs1513 genotypes (recessive effect).

| Model | Statistic | Fixed effects |  |  | Random effects |  |  |
| --- | --- | --- | --- | --- | --- | --- | --- |
|  |  | FADS2 |  | G×E | FADS2 |  | G×E |
|  |  | Other genotypes | GG |  | Other genotypes | GG |  |
| rs174575 (CC or CG=0; GG=1) |  |  |  |  |  |  |  |
| Unadjusted | $I^2$ | - | - | - | 76.4 | 64.4 | 77.6 |
| $N_{\text{estimates}}=8$ | P-value | $8.6\times10^{-50}$ | $3.8\times10^{-8}$ | 0.188 | $7.6\times10^{-7}$ | 0.002 | 0.323 |
| $N_{\text{subjects}}=12,614$ | $\beta$ | 0.37 | 0.43 | 0.11 | 0.29 | 0.43 | 0.18 |
|  | 95% CI | 0.32; 0.41 | 0.28; 0.58 | -0.05; 0.27 | 0.17; 0.40 | 0.16; 0.70 | -0.18; 0.54 |
| Adjusted (1)<sup>a</sup> | $I^2$ | - | - | - | 74.2 | 67.2 | 75.5 |
| $N_{\text{estimates}}=8$ | P-value | $7.7\times10^{-48}$ | $9.3\times10^{-7}$ | 0.603 | $7.9\times10^{-7}$ | 0.024 | 0.705 |
| $N_{\text{subjects}}=12,590$ | $\beta$ | 0.37 | 0.39 | 0.04 | 0.29 | 0.35 | 0.07 |
|  | 95% CI | 0.32; 0.42 | 0.23; 0.54 | -0.12; 0.21 | 0.18; 0.41 | 0.04; 0.65 | -0.29; 0.43 |
| Adjusted (2)<sup>b</sup> | $I^2$ | - | - | - | 84.1 | 47.4 | 59.5 |
| $N_{\text{estimates}}=8$ | P-value | $6.4\times10^{-20}$ | $6.4\times10^{-5}$ | 0.244 | 0.055 | 0.020 | 0.445 |
| $N_{\text{subjects}}=12,077$ | $\beta$ | 0.25 | 0.34 | 0.10 | 0.15 | 0.31 | 0.12 |
|  | 95% CI | 0.20; 0.31 | 0.17; 0.51 | -0.07; 0.28 | 0.00; 0.31 | 0.05; 0.58 | -0.19; 0.43 |
| rs1535 (AA or AG=0; GG=1) |  |  |  |  |  |  |  |
| Unadjusted | $I^2$ | - | - | - | 73.5 | 54.1 | 42.6 |
| $N_{\text{estimates}}=9$ | P-value | $9.2\times10^{-49}$ | $2.2\times10^{-6}$ | 0.663 | $4.6\times10^{-7}$ | 0.013 | 0.646 |
| $N_{\text{subjects}}=13,202$ | $\beta$ | 0.37 | 0.29 | -0.03 | 0.29 | 0.24 | -0.04 |
|  | 95% CI | 0.32; 0.42 | 0.17; 0.41 | -0.16; 0.10 | 0.18; 0.40 | 0.05; 0.43 | -0.24; 0.15 |
| Adjusted (1)<sup>a</sup> | $I^2$ | - | - | - | 76.0 | 47.7 | 60.9 |
| $N_{\text{estimates}}=9$ | P-value | $9.9\times10^{-47}$ | $2.2\times10^{-7}$ | 0.720 | $7.1\times10^{-6}$ | $5.4\times10^{-3}$ | 0.778 |
| $N_{\text{subjects}}=13,175$ | $\beta$ | 0.37 | 0.33 | -0.02 | 0.29 | 0.27 | -0.03 |
|  | 95% CI | 0.32; 0.42 | 0.20; 0.45 | -0.16; 0.11 | 0.16; 0.42 | 0.08; 0.47 | -0.28; 0.21 |
| Adjusted (2)<sup>b</sup> | $I^2$ | - | - | - | 84.0 | 25.9 | 49.6 |
| $N_{\text{estimates}}=9$ | P-value | $1.9\times10^{-19}$ | $1.2\times10^{-5}$ | 0.277 | 0.065 | 0.003 | 0.592 |
| $N_{\text{subjects}}=12,633$ | $\beta$ | 0.26 | 0.28 | 0.07 | 0.15 | 0.25 | 0.06 |
|  | 95% CI | 0.20; 0.31 | 0.16; 0.41 | -0.06; 0.21 | -0.01; 0.32 | 0.09; 0.41 | -0.16; 0.27 |

<sup>a</sup>Covariates were sex, age (linear and quadratic terms), ancestry-informative principal components (if available) and genotyping centre (if necessary).

a Covariates were sex, age (linear and quadraric terms), ancestry-informative principal components (if available) and genotypingcentre (if necessary).

b Same covariates than in the Adjusted (1) model, in addition to maternal education (linear and quadratic terms) and/or maternal cognition (linear and quadratic terms). GxE: interaction between breastfeeding and polymorphisms in the *FADS2* gene. N_estimates_: number of estimates being pooled. N_subjects_: pooled sample size.

**Figure 1.**
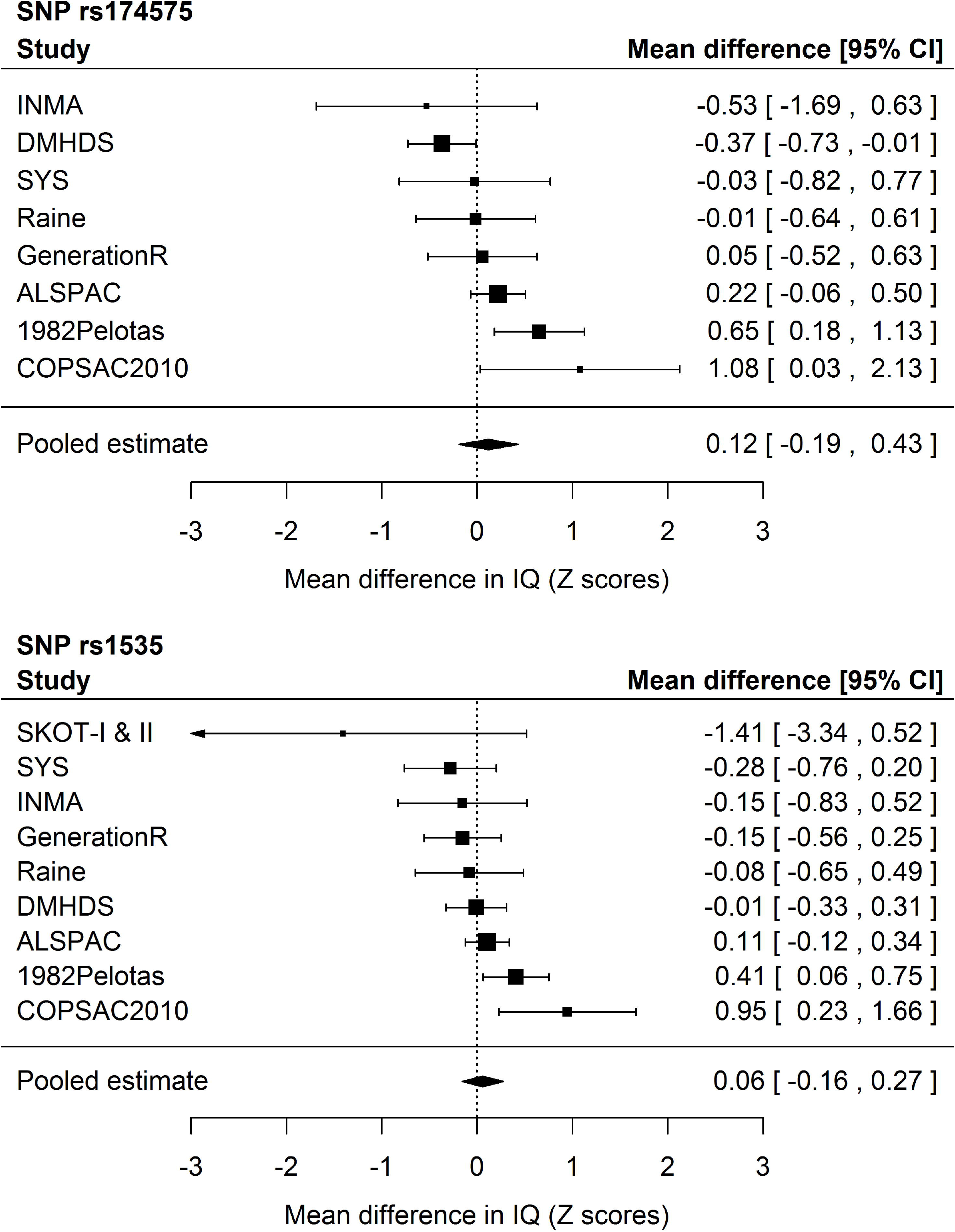
Forest plots of mean differences in IQ Z scores from the fully adjusted model comparing ever with never breastfed individuals based on random effects meta-analysis. SKOT-I and SKOT-II were excluded from the analyses for the rs174575 polymorphism because the model did not fit (due to a combination of modest sample size, high prevalence of breastfeeding and assuming a recessive genetic effect of the rarest allele). 1982Pelotas: 1982 Pelotas Birth Cohort. ALSPAC: Avon Longitudinal Study of Parents and Children. COPSAC2010: Copenhagen Prospective Study on Asthma in Childhood 2010. DMHDS: Dunedin Multidisciplinary Health and Development Study. GenerationR: Generation R Study. INMA: *INfancia y Medio Ambiente* - Environment and Childhood. Raine: Western Australian Pregnancy Cohort (Raine) Study. SKOT-I & II: *Småbørn Kost Og Trivsel* (I and II). SYS: Saguenay Youth Study.

### Secondary analysis

As shown in Table 2 and Supplementary Tables 4-6, there was no strong indication of interaction when analysing other categorisations of breastfeeding duration and *FADS2* SNPs coded assuming a recessive effect. This was also the case when *FADS2* variants were coded assuming additive (Supplementary Table 7), dominant (Supplementary Table 8) and overdominant (Supplementary Table 9) effects. The same was observed for exclusive breastfeeding (Supplementary Tables 10-13).

**Table 2.** Meta-analytical linear regression coefficients (β) of the interaction term between *FADS2* rs174575 or rs1535 genotypes (recessive effect) with breastfeeding (<6 months vs. >6 months, in ordinal categories or in months), having cognitive measures (in standard deviation units) as the outcome.

| Model | Statistic | Fixed effects |  |  | Random effects |  |  |
| --- | --- | --- | --- | --- | --- | --- | --- |
|  |  | <6 months=0<br>≥6 months=1 | Numerically-<br>coded<br>categories | Months | <6 months=0<br>≥6 months=1 | Numerically-<br>coded<br>categories | Months |
| rs174575 (CC or CG=0; GG=1) |  |  |  |  |  |  |  |
| Unadjusted | $I^2$ | - | - | - | 23.1 | 57.1 | 13.9 |
| $N_{\text{estimates}}=8$ | P-value | 0.515 | 0.104 | 0.371 | 0.647 | 0.150 | 0.335 |
| $N_{\text{subjects}}=11,733$ | $\beta$ | 0.05 | 0.04 | 0.01 | 0.04 | 0.06 | 0.01 |
|  | 95% CI | -0.10; 0.20 | -0.01; 0.09 | -0.01; 0.02 | -0.14; 0.22 | -0.02; 0.15 | -0.01; 0.03 |
| Adjusted (1)<sup>a</sup> | $I^2$ | - | - | - | 53.6 | 58.7 | 63.3 |
| $N_{\text{estimates}}=8$ | P-value | 0.378 | 0.189 | 0.608 | 0.546 | 0.282 | 0.635 |
| $N_{\text{subjects}}=11,706$ | $\beta$ | 0.07 | 0.04 | 0.00 | 0.08 | 0.06 | 0.01 |
|  | 95% CI | -0.09; 0.23 | -0.02; 0.09 | -0.01; 0.02 | -0.18; 0.35 | -0.05; 0.16 | -0.02; 0.04 |
| Adjusted (2)<sup>b</sup> | $I^2$ | - | - | - | 82.6 | 84.6 | 85.3 |
| $N_{\text{estimates}}=8$ | P-value | 0.244 | 0.132 | 0.782 | 0.496 | 0.346 | 0.602 |
| $N_{\text{subjects}}=11,242$ | $\beta$ | 0.10 | 0.04 | 0.00 | 0.17 | 0.09 | 0.01 |
|  | 95% CI | -0.07; 0.26 | -0.01; 0.10 | -0.01; 0.02 | -0.32; 0.65 | -0.09; 0.26 | -0.04; 0.07 |
| rs1535 (AA or AG=0; GG=1) |  |  |  |  |  |  |  |
| Unadjusted | $I^2$ | - | - | - | 0.0 | 0.0 | 0.0 |
| $N_{\text{estimates}}=8$ | P-value | 0.460 | 0.966 | 0.805 | 0.460 | 0.966 | 0.805 |
| $N_{\text{subjects}}=12,018$ | $\beta$ | -0.05 | 0.00 | 0.00 | -0.05 | 0.00 | 0.00 |
|  | 95% CI | -0.17; 0.08 | -0.04; 0.04 | -0.01; 0.01 | -0.17; 0.08 | -0.04; 0.04 | -0.01; 0.01 |
| Adjusted (1)<sup>a</sup> | $I^2$ | - | - | - | 8.0 | 54.3 | 59.6 |
| $N_{\text{estimates}}=8$ | P-value | 0.248 | 0.508 | 0.538 | 0.302 | 0.635 | 0.330 |
| $N_{\text{subjects}}=11,991$ | $\beta$ | -0.07 | -0.01 | 0.00 | -0.07 | -0.02 | -0.01 |
|  | 95% CI | -0.20; 0.05 | -0.06; 0.03 | -0.01; 0.01 | -0.20; 0.06 | -0.09; 0.05 | -0.03; 0.01 |
| Adjusted (2)<sup>b</sup> | $I^2$ | - | - | - | 3.9 | 29.9 | 35.5 |
| $N_{\text{estimates}}=8$ | P-value | 0.194 | 0.675 | 0.320 | 0.216 | 0.728 | 0.344 |
| $N_{\text{subjects}}=11,499$ | $\beta$ | -0.08 | -0.01 | -0.01 | -0.08 | -0.01 | -0.01 |
|  | 95% CI | -0.21; 0.04 | -0.05; 0.03 | -0.02; 0.01 | -0.21; 0.05 | -0.07; 0.05 | -0.02; 0.01 |

a Covariates were sex, age (linear and quadratic terms), ancestry-informative principal components (if available) and genotyping centre (if necessary).

b Same covariates than in the Adjusted (1) model, in addition to maternal education (linear and quadratic terms) and/or maternal cognition (linear and quadratic terms). N_estimates_: number of estimates being pooled. N_subjects_: pooled sample size.

Supplementary Table 14 displays the results obtained when including the UK Biobank, which was analysed as two independent samples according to the genotyping platform (Biobank_Axiom and Biobank_BiLEVE). Its inclusion resulted in a combined sample size of more than 45,000 individuals. When *FADS2* variants were coded assuming recessive effects, the pooled estimates from the unadjusted model −0.02 (95% CI: −0.10; 0.06) and 0.08 (95% CI: −0.13; 0.29) for fixed and random-effects meta-analysis, respectively. The corresponding estimates from the adjusted (1) model were −0.04 (95% CI: −0.13; 0.04) and 0.00 (95% CI: −0.21; 0.20), respectively. There was also no strong statistical evidence supporting an interaction when other genetic effects were assumed.

### Sensitivity analysis

Table 3 displays the results of random-effects meta-regression. Neither type of IQ test, timing of breastfeeding measurement, continental region nor mean year of birth explained a substantial amount of between-study heterogeneity. For rs174575, the adjusted R^2^ of ancestry-informative principal components was 88.0%, with pooled estimates of 0.28 (95% CI: 0.02; 0.54) and −0.38 (95% CI: −0.72; −0.04) Z scores in IQ from studies that did and did not adjust for principal components, respectively, which would be suggestive of confounding due to population stratification towards a negative association. Age at IQ measurement was inversely associated with the magnitude of the interaction term, with pooled estimates of 0.06 (95% CI: −0.46; 0.58) and 0.20 (95% CI: −0.18; 0.58) when IQ was measured at 10 years of age or more, or before that age (respectively), possibly suggesting an attenuation of the effect over time. The adjusted R^2^ was 10.4% when entering age as a continuous variable, but 0% when dichotomised. When stratifying studies according to prevalence of ever breastfeeding, the pooled estimate among studies with a prevalence ≥90% was 0.36 (95% CI: −0.19; 0.90), and −0.04 (95% CI: −0.38; 0.29) when pooling the remaining studies. Adjusted R^2^ estimates were 16.4% and 72.3% when prevalence of ever breastfeeding was analysed as a binary and as a continuous variable, respectively. Among studies with breastfeeding duration equal to or greater than the median among studies (i.e., 5.85 months), the pooled estimate was 0.50 (95% CI: −0.06; 1.06), compared to 0.14 (95% CI: −0.10; 0.38) when pooling the remaining studies. The adjusted R^2^ was 45.5% when breastfeeding duration was dichotomised at the median, but 0% when analysed continuously. When stratifying studies into larger (≥1000 individuals) and smaller (<1000 individuals), the pooled estimates were 0.26 (95% CI: 0.00; 0.52) and −0.03 (95% CI: −0.63; 0.56), with an adjusted R^2^ of 33.8% when sample size was dichotomised, and of 0% when analysed in continuous form.

**Table 3.**
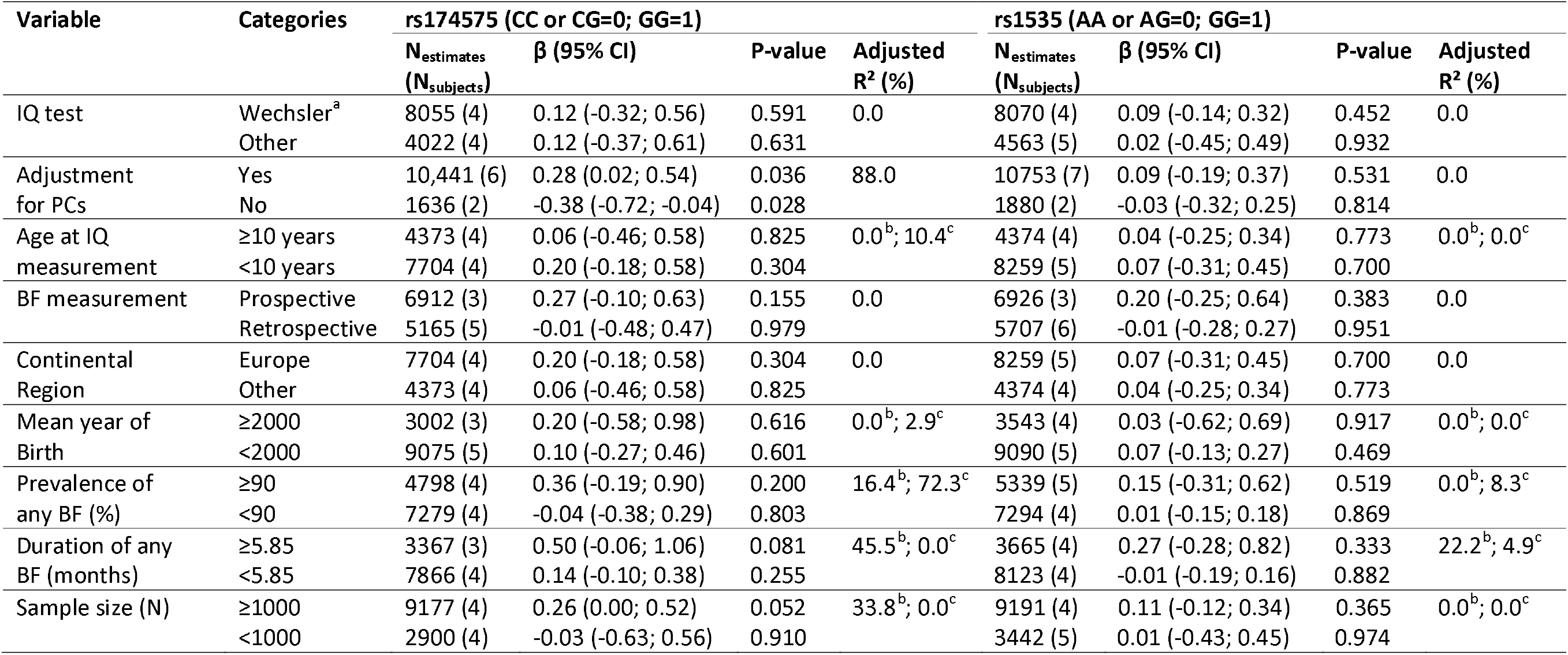
Stratified random effects meta-analytical linear regression coefficients (β) of the interaction term between *FADS2* rs174575 or rs1535 genotypes (recessive effect) with breastfeeding (never=0; ever=l), having cognitive measures (in standard deviation units) as the outcome. Estimates from the fully adjusted model were used.

| Variable | Categories | rs174575 (CC or CG=0; GG=1) |  |  |  | rs1535 (AA or AG=0; GG=1) |  |  |  |
| --- | --- | --- | --- | --- | --- | --- | --- | --- | --- |
| | | N<sub>estimates</sub><br>(N<sub>subjects</sub>) | $\beta$ (95% CI) | P-value | Adjusted<br>R<sup>2</sup> (%) | N<sub>estimates</sub><br>(N<sub>subjects</sub>) | $\beta$ (95% CI) | P-value | Adjusted<br>R<sup>2</sup> (%) |
| IQ test | Wechsler<sup>a</sup> | 8055 (4) | 0.12 (-0.32; 0.56) | 0.591 | 0.0 | 8070 (4) | 0.09 (-0.14; 0.32) | 0.452 | 0.0 |
|  | Other | 4022 (4) | 0.12 (-0.37; 0.61) | 0.631 |  | 4563 (5) | 0.02 (-0.45; 0.49) | 0.932 |  |
| Adjustment<br>for PCs | Yes | 10,441 (6) | 0.28 (0.02; 0.54) | 0.036 | 88.0 | 10753 (7) | 0.09 (-0.19; 0.37) | 0.531 | 0.0 |
|  | No | 1636 (2) | -0.38 (-0.72; -0.04) | 0.028 |  | 1880 (2) | -0.03 (-0.32; 0.25) | 0.814 |  |
| Age at IQ<br>measurement | ≥10 years | 4373 (4) | 0.06 (-0.46; 0.58) | 0.825 | 0.0<sup>b</sup>; 10.4<sup>c</sup> | 4374 (4) | 0.04 (-0.25; 0.34) | 0.773 | 0.0<sup>b</sup>; 0.0<sup>c</sup> |
|  | <10 years | 7704 (4) | 0.20 (-0.18; 0.58) | 0.304 |  | 8259 (5) | 0.07 (-0.31; 0.45) | 0.700 |  |
| BF measurement | Prospective | 6912 (3) | 0.27 (-0.10; 0.63) | 0.155 | 0.0 | 6926 (3) | 0.20 (-0.25; 0.64) | 0.383 | 0.0 |
|  | Retrospective | 5165 (5) | -0.01 (-0.48; 0.47) | 0.979 |  | 5707 (6) | -0.01 (-0.28; 0.27) | 0.951 |  |
| Continental<br>Region | Europe | 7704 (4) | 0.20 (-0.18; 0.58) | 0.304 | 0.0 | 8259 (5) | 0.07 (-0.31; 0.45) | 0.700 | 0.0 |
|  | Other | 4373 (4) | 0.06 (-0.46; 0.58) | 0.825 |  | 4374 (4) | 0.04 (-0.25; 0.34) | 0.773 |  |
| Mean year of<br>Birth | ≥2000 | 3002 (3) | 0.20 (-0.58; 0.98) | 0.616 | 0.0<sup>b</sup>; 2.9<sup>c</sup> | 3543 (4) | 0.03 (-0.62; 0.69) | 0.917 | 0.0<sup>b</sup>; 0.0<sup>c</sup> |
|  | <2000 | 9075 (5) | 0.10 (-0.27; 0.46) | 0.601 |  | 9090 (5) | 0.07 (-0.13; 0.27) | 0.469 |  |
| Prevalence of<br>any BF (%) | ≥90 | 4798 (4) | 0.36 (-0.19; 0.90) | 0.200 | 16.4<sup>b</sup>; 72.3<sup>c</sup> | 5339 (5) | 0.15 (-0.31; 0.62) | 0.519 | 0.0<sup>b</sup>; 8.3<sup>c</sup> |
|  | <90 | 7279 (4) | -0.04 (-0.38; 0.29) | 0.803 |  | 7294 (4) | 0.01 (-0.15; 0.18) | 0.869 |  |
| Duration of any<br>BF (months) | ≥5.85 | 3367 (3) | 0.50 (-0.06; 1.06) | 0.081 | 45.5<sup>b</sup>; 0.0<sup>c</sup> | 3665 (4) | 0.27 (-0.28; 0.82) | 0.333 | 22.2<sup>b</sup>; 4.9<sup>c</sup> |
|  | <5.85 | 7866 (4) | 0.14 (-0.10; 0.38) | 0.255 |  | 8123 (4) | -0.01 (-0.19; 0.16) | 0.882 |  |
| Sample size (N) | ≥1000 | 9177 (4) | 0.26 (0.00; 0.52) | 0.052 | 33.8<sup>b</sup>; 0.0<sup>c</sup> | 9191 (4) | 0.11 (-0.12; 0.34) | 0.365 | 0.0<sup>b</sup>; 0.0<sup>c</sup> |
|  | <1000 | 2900 (4) | -0.03 (-0.63; 0.56) | 0.910 |  | 3442 (5) | 0.01 (-0.43; 0.45) | 0.974 |  |

<sup>b</sup>Variable categorised as shown in the table.

a Includes both Wechsler Adult Intelligence Scale (ALSPAC and Dunedin Multidisciplinary Health and Development Study) and Wechsler Intelligence Scale for Children (1982 Pelotas Birth Cohort and Saguenay Youth Study).

b Variable categorised as shown in the table.

c Variable entered in continuous form (e.g., age at outcome measurement modelled in years, as a continuous variable). PCs: ancestry-informative genetic principal components. BF: breastfeeding. N: number of. CI: confidence interval N_estimates_: number of estimates being pooled. N_subjects_: pooled sample size.

Regarding the rs1535 variant, the respective subgroup-specific estimates were consistent with those of the rs174575 SNP: adjustment for principal components, with pooled estimates of 0.09 (95% CI: −0.19; 0.37) and −0.03 (95% CI: −0.32; 0.25) among studies that did and did not perform this adjustment, respectively; age at IQ measurement, with pooled estimates of 0.04 (95% CI: −0.19; 0.37) and 0.07 (95% CI: −0.31; 0.45) among studies that measured IQ when individuals were ≥10 and <10 years-old, respectively; and sample size, with pooled estimates of 0.11 (95% CI: −0.12; 0.34) and 0.01 (95% CI: −0.43 and 0.45) among larger and smaller studies, respectively. However, in all those cases the adjusted R^2^ values were 0%. Prevalence of ever breastfeeding presented adjusted R^2^ values of 0% and 8.3% when dichotomised and analysed continuously, respectively. The pooled estimates for the rs1535 variant were 0.15 (95% CI: −0.31; 0.62) and 0.01 (95% CI: −0.15; 0.18) among studies with prevalence of ever breastfeeding of ≥90% and <90%, respectively. The most consistent moderator between SNPs was breastfeeding duration, with pooled estimates for the rs1535 SNP of 0.27 (95% CI: −0.28; 0. 82) and −0.01 (95% CI: −0.19; 0.16) among studies with ≥5.85 and <5.85 months of duration, respectively; adjusted R^2^ values were 22.2% and 4.9% when breastfeeding duration was dichotomised and analysed continuously, respectively (Figure 2).

**Figure 2.**
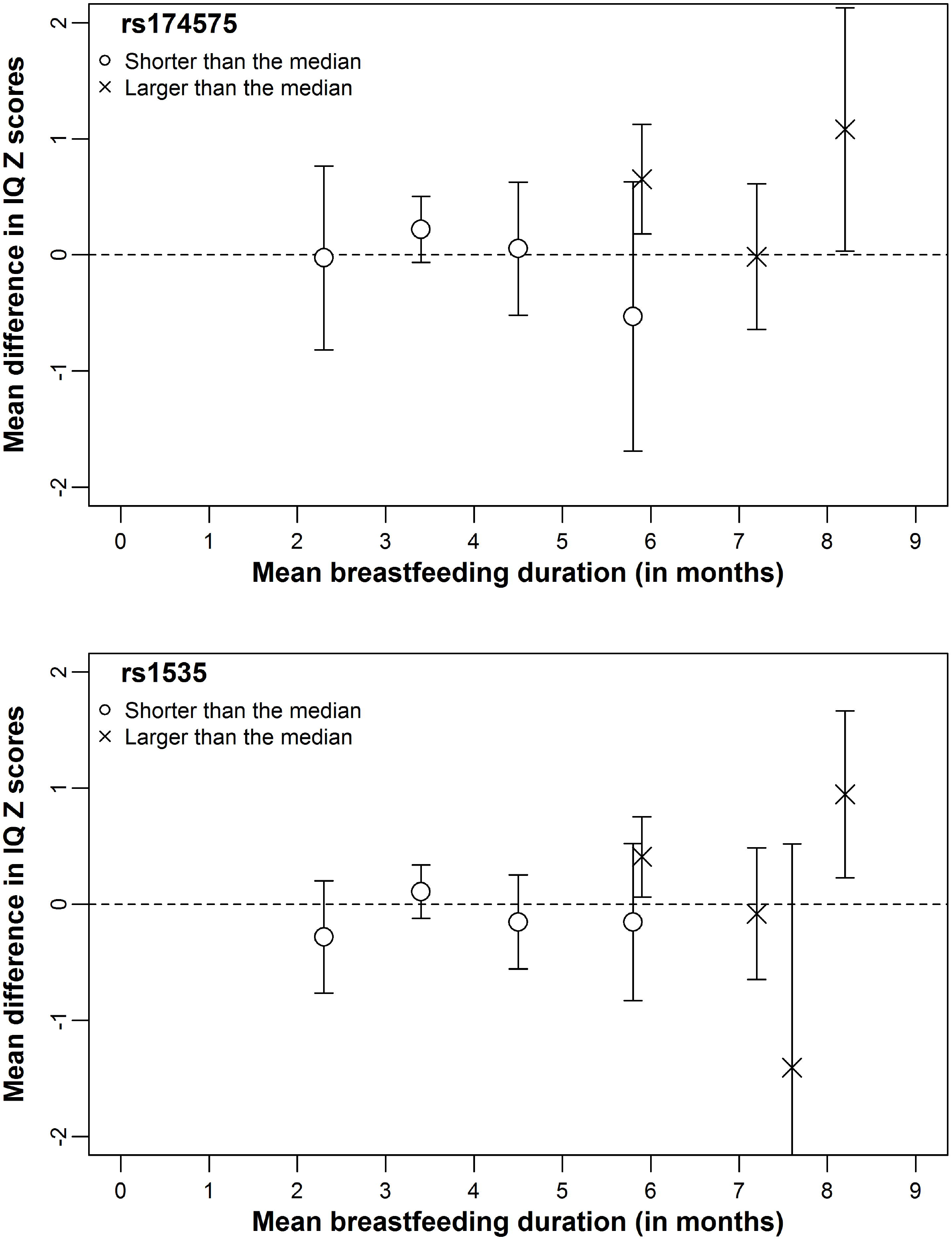
Scatter plots of mean differences (with 95% confidence intervals) in IQ Z scores from the fully adjusted model comparing ever with never breastfed individuals according to prevalence (%) of ever breastfeeding and average breastfeeding duration in months.

There was no strong evidence in support of gene-environment correlation involving maternal education or maternal cognition (Table 4). Regarding the rs174575 variant, random effects meta-analytical estimates from the adjusted model were 0.16 (95% CI: −0.45; 0.78) for maternal education, and −0.02 (95% CI: −0.25; 0.21) for maternal cognition, respectively. The corresponding estimates for the rs1535 SNP were −0.12 (95% CI: −0.51; 0.27) and 0.14 (95% CI: −0.04; 0.33).

**Table 4.** Meta-analytical linear regression coefficients (β) of the interaction term between *FADS2* rs174575 or rs1535 genotypes (recessive effect) with breastfeeding (never=0; ever=l), having maternal education (in complete years) or maternal cognitive measures (in standard deviation units) as the outcome.

| Model | Statistic | Fixed effects |  | Random effects |  |
| --- | --- | --- | --- | --- | --- |
|  |  | Maternal education | Maternal cognition | Maternal education | Maternal cognition |
| rs174575 (CC or CG=0; GG=1) |  |  |  |  |  |
| Unadjusted | N<sub>estimates</sub> | 7 | 5 | 7 | 5 |
|  | N<sub>subjects</sub> | 14,671 | 6299 | 14671 | 6299 |
|  | I<sup>2</sup> | - | - | 81.1 | 18.1 |
|  | P-value | 0.159 | 0.326 | 0.375 | 0.389 |
|  | β | 0.28 | 0.10 | 0.59 | 0.10 |
|  | 95% CI | -0.11; 0.66 | -0.10; 0.31 | -0.72; 1.91 | -0.13; 0.33 |
| Adjusted (1)<sup>a</sup> | N<sub>estimates</sub> | 7 | 5 | 7 | 5 |
|  | N<sub>subjects</sub> | 12,113 | 6126 | 12113 | 6126 |
|  | I<sup>2</sup> | - | - | 14.1 | 0.0 |
|  | P-value | 0.509 | 0.854 | 0.607 | 0.854 |
|  | β | 0.16 | -0.02 | 0.16 | -0.02 |
|  | 95% CI | -0.31; 0.62 | -0.25; 0.21 | -0.45; 0.78 | -0.25; 0.21 |
| rs1535 (AA or AG=0; GG=1) |  |  |  |  |  |
| Unadjusted | N<sub>estimates</sub> | 8 | 5 | 8 | 5 |
|  | N<sub>subjects</sub> | 15,447 | 6556 | 15447 | 6556 |
|  | I<sup>2</sup> | - | - | 1.4 | 0.0 |
|  | P-value | 0.784 | 0.272 | 0.814 | 0.272 |
|  | β | -0.05 | 0.10 | -0.04 | 0.10 |
|  | 95% CI | -0.38; 0.28 | -0.08; 0.28 | -0.39; 0.31 | -0.08; 0.28 |
| Adjusted (1)<sup>a</sup> | N<sub>estimates</sub> | 8 | 5 | 8 | 5 |
|  | N<sub>subjects</sub> | 12,743 | 6378 | 12743 | 6378 |
|  | I<sup>2</sup> | - | - | 0.0 | 0.0 |
|  | P-value | 0.540 | 0.160 | 0.540 | 0.160 |
|  | β | -0.12 | 0.14 | -0.12 | 0.14 |
|  | 95% CI | -0.51; 0.27 | -0.05; 0.33 | -0.51; 0.27 | -0.05; 0.33 |

<sup>a</sup>Covariates were sex, age (linear and quadratic terms), ancestry-informative principal components (if available) and genotyping centre (if necessary).

a Covariates were sex, age (linear and quatric terms), ancestry-informative principal components (if available) and genotyping centre (if necessary). N_estimates_: number of estimates being pooled. N_subjects_: pooled sample size.

## Discussion

Our primary analyses were not supportive of the hypothesis that the *FADS2* polymorphisms rs174575 and rs1535 and breastfeeding interact to affect IQ. This was also the case in a priori secondary analyses using different categorisations of breastfeeding, exclusive rather than any quality of breastfeeding, assuming different genetic effects and including a large study that did not meet all eligibility criteria. Sensitivity analyses were not supportive that gene-environment correlation involving maternal education or maternal cognition substantially influenced the results. Random effects meta-regression suggested that breastfeeding duration was an important moderator.

Results from our primary and secondary analyses were not supportive of the nutritional adequacy hypothesis, according to which a positive interaction coefficient would be expected^14^. In other words, there might be no upper limit (or it may be very high) of the effects of LC-PUFAs on IQ, so that supplementing infants with LC-PUFAs could be beneficial for cognition for both lactating and non-lactating infants alike. Importantly, this does not imply that LC-PUFAs supplementation completely replaces the benefits of breastfeeding, since the latter may act through diverse mechanisms, and also provide benefits other than for intelligence^1,47^.

On the other hand, in our random effects meta-regression analysis, studies with longer average breastfeeding duration generally presented interaction coefficients that were positive and stronger in magnitude than studies with shorter breastfeeding duration. Moreover, average breastfeeding duration was the most consistent moderator between polymorphisms. Considering that positive interaction coefficients are expected under the nutritional adequacy hypothesis, this result raises the possibility that there may be an upper limit of the benefits of LC-PUFAs, but achieving such limits from breast milk requires that breastfeeding lasts for some (currently unknown) time. Given that breastfeeding practices in the participating studies were generally well below international recommendations^48,49^, it is possible that the amount of LC-PUFA received from breast milk were, on average, lower than this threshold.

The strengths of our study include: appropriate sample size for the primary analysis^26^; publication of study protocol^26^, which helps to avoid biased reporting; analyses performed using standardised analysis scripts and harmonised (as much as possible) datasets; inclusion of published and unpublished reports, thus minimising publication bias; several *a priori* defined secondary and sensitivity analyses; proper adjustment for covariates in the G×E setting; and IQ measures with similar variances, which reduces heterogeneity that could arise due to Z score conversion^50,51^.

Our study also had limitations. Some of them were related to the small numbers of individuals in some categories, which we tried to resolve by changing the protocol, such as in the case of the definition of never being breastfed and exclusion of some categorisations of breastfeeding from the analysis. Indeed, had the latter been maintained, the hypothesis above regarding breastfeeding duration and nutritional adequacy could have been studied. However, due to statistical issues, we opted for excluding this variable. Other limitations were: small sample size for some analyses, such as those involving exclusive breastfeeding; heterogeneity in important study characteristics, such as age, IQ test, timing of breastfeeding measurement, etc.; and small number of studies for meta-regression analyses. Another potential limitation is lack of adjustment for maternal genotypes, which may confound the association between participant’s genotype and IQ by influencing fatty acid composition in breast milk^25^. However, although there is evidence that this may be the case for some genetic variants implicated in LC-PUFA metabolism^25^, there is no strong evidence that maternal genotypes with regards to the particular SNPs that we studied are associated with offspring’s IQ or that they interact with breastfeeding^14^. It is also possible that there are epistatic relationships between genes implicated in this pathway, so that focusing only on two variants in a single gene may not capture the whole complexity of the interplay between genetic influences in LC-PUFA levels, breastfeeding and cognitive development.

Although our primary findings were not supportive of an interaction between breastfeeding and *FADS2* polymorphisms, random effects meta-regression results suggest that such interaction exist, with studies with longer average breastfeeding duration generally presenting estimates in accordance with the nutritional adequacy hypothesis. This should be investigated in future studies comparing different categories of breastfeeding duration, rather than simply never vs. ever comparisons (or other categorisations used here). Since such analysis would involve many subgroupings, the best alternative is likely to perform such analysis in a large dataset of individual-level data, which may be achieved by a consortium-based effort such as this collaborative meta-analysis. This and other future investigations will be important to further refine our understanding on the role of LC-PUFAs on the association between breastfeeding and intelligence. This will also have more practical implications, such as identifying whether current breastfeeding recommendations allow achieving the upper limit of cognitive benefits related to LC-PUFAs intake (if such limit exists), and the potential benefits (if any) of supplementing a lactating infant with LC-PUFAs.

